# The what and where of synchronous sound perception

**DOI:** 10.1101/2021.12.22.473782

**Authors:** Guus C. Van Bentum, A. John Van Opstal, Marc M. Van Wanrooij

## Abstract

Sound localization and identification are challenging in acoustically rich environments. The relation between these two processes is still poorly understood. As natural sound-sources rarely occur exactly simultaneously, we wondered whether the auditory system could identify (“what”) and localize (“where”) two spatially separated sounds with synchronous onsets. While listeners typically report hearing a single source at an average location, one study found that both sounds may be accurately localized if listeners are explicitly being told two sources exist. We here tested whether simultaneous source identification (one vs. two) and localization is possible, by letting listeners choose to make either one or two head-orienting saccades to the perceived location(s). Results show that listeners could identify two sounds only when presented on different sides of the head, and that identification accuracy increased with their spatial separation. Notably, listeners were unable to accurately localize either sound, irrespective of whether one or two sounds were identified. Instead, the first (or only) response always landed near the average location, while second responses were unrelated to the targets. We conclude that localization of synchronous sounds in the absence of prior information is impossible. We discuss that the putative cortical ‘what’ pathway may not transmit relevant information to the ‘where’ pathway. We examine how a broadband interaural correlation cue could help to correctly identify the presence of two sounds without being able to localize them. We propose that the persistent averaging behavior reveals that the ‘where’ system intrinsically assumes that synchronous sounds originate from a single source.

**Significance Statement:** It is poorly understood whether identification (‘what’) of sounds and their localization (‘where’) are inter-related, or independent neural processes. We measured sound-localization responses towards synchronous sounds to examine potential coupling of these processes. We varied the spatial configurations of two sounds and found that although identification improved considerably with larger spatial separation, their localization was unaffected: responses were always directed towards the average location. This shows absence of mutual coupling of information between the ‘what’ and ‘where’ streams in the auditory system. We also show how broadband interaural correlation could explain the improved identification results, without affecting localization performance, and explain how the persistent spatial averaging could be understood from strong internal priors regarding sound synchronicity.

## Introduction

There is ample evidence that in the visual system an object’s identity (‘what’) and its location (‘where’) are analyzed by separate processing streams, known as the ventral and dorsal neural pathways, respectively (Goodale and Milner, 1991). Likewise, the auditory system is faced with the problem to identify and localize sounds in the presence of background noise, competing stimuli, or acoustic reflections (Maeder, 2001). Because the acoustic input is tonotopically, rather than spatially, organized this is a challenging task, especially when spectral-temporal acoustic signals overlap. Caudal and rostral pathways in auditory cortex could fulfill similar functions (Romanski et al., 1999; Bizley and Cohen, 2013). Electrophysiological studies in non-human primates implicate that rostral auditory-cortical areas primarily process sound-object recognition and identity, whereas spatial processing predominantly occurs in caudal auditory areas (Recanzone and Cohen, 2010; Rauschecker and Tian, 2000). Lesions (Clarke et al., 2002) and fMRI/ERP studies (Alain et al., 2001; Ahveninen et al., 2006) further support the idea of separated pathways for source identification and localization in the human auditory cortex.

Obviously, the auditory system can readily perform sound-source identification, segregation, and localization when sources differ substantially in their spectral-temporal content, and when they are widely separated in space (Best et al., 2004) and time (Brown et al., 2015; Ege et al., 2018). However, for synchronous sounds, spectral-temporal overlap may pose considerable problems for source identification and localization. Typically, identification (Litovsky and Shinn-Cunningham, 2001) and localization (in azimuth: Blauert, 1971; Perrot et al., 1987; Van Bentum et al., 2021; in elevation: Bremen et al., 2010; Brown et al., 2015; Van Bentum et al., 2017; Ege et al., 2018) of synchronous sounds are both (Seeber and Hafner, 2011) inaccurate; listeners report hearing a single sound at the average location of the two sound sources. Yet, when listeners are forced to localize two synchronous sounds in the horizontal plane in repeated trials, accurate performance is possible (Yost and Brown, 2013). This may suggest that even though an accurate representation of both locations is available to the auditory-localization system, an average localization response is programmed. In none of these studies, however, source identification and localization were tested simultaneously, so that it remains unclear whether they share the same, or use independent, neural mechanisms. To reveal their potential interaction requires tests in which localization and identification of a pair of synchronous sounds are assessed within the same trial and within the same paradigm.

Here, normal-hearing listeners made goal-directed head movements to two synchronous, uncorrelated broadband noise bursts, which were perceptually identical. Listeners had no prior information regarding the number of sounds and were instructed to make either one or two goal-directed responses to indicate number and location(s) of the perceived sound(s). Their head movements thus served as proxies for source-identification and localization. We systematically varied spatial separation of the sounds and presented them either on the same side of the head (within-hemisphere), or on different sides (cross-hemisphere). This manipulation effectively created sound pairs with higher or lower interaural correlation, respectively, which could potentially aid in the segregation of acoustic events (Faller and Merimaa, 2004; May et al., 2013). This allowed us to study use of this potential segregation cue by the putative localization and identification pathways, and their potential mutual coupling (Fig. 1). If these pathways are coupled, listeners who identified two sounds will also be able to localize them. If they are uncoupled, however, these outcomes will be unrelated.

**Figure 1.**
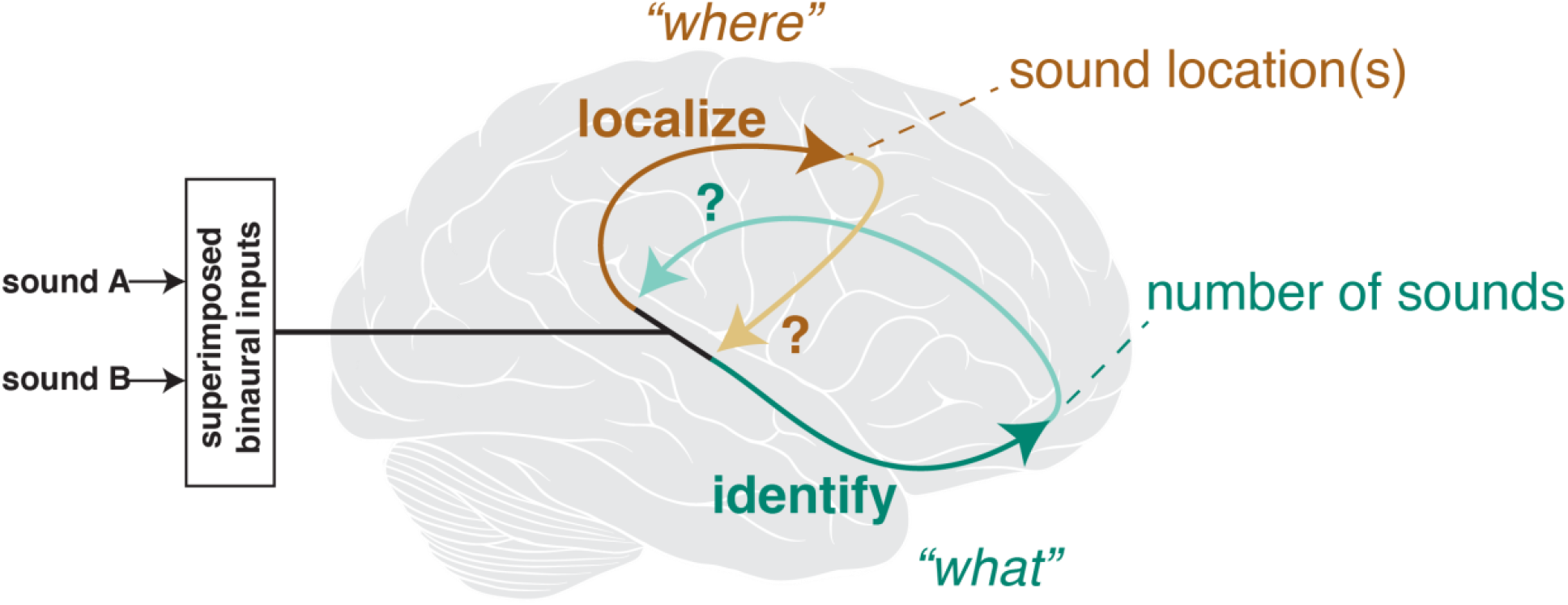
Rationale. Two conceptual schemes that differ as to whether sound-source identification (successful detection of one or two sources) and localization use coupled or independent neural mechanisms. Listeners responded to the percept of either one or two synchronous sound sources (A and B). In case of coupled ‘what’ and ‘where’ streams, accurate identification (green arrows) could signal information to the localization stream, and vice versa (brown arrows). In case of uncoupled systems, these mutual projections are absent, and successful source identification will be unrelated to localization performance.

## Methods

### Listeners

Six adult, normal-hearing listeners (three male, ages 22-32) participated in this study. Three listeners had no prior experience with sound-localization experiments in the laboratory. All listeners, except for the first author, were naïve regarding the purpose of the experiment. All experimental procedures were approved by the local Ethics Committee of the Faculty of Social Sciences of Radboud University (protocol nr. ECSW2016-2208-41). Subjects gave their full written informed consent, prior to their participation in these experiments. Listeners took part in a larger study containing nine experimental sessions on different days, in which task instructions and the percentage of trials with one or two sounds were varied. Here, we present the results from one session, which featured *only* trials with two sounds. As subjects were not aware of this (experimental sessions were randomized) they had no prior knowledge about the number of sound sources in each trial, nor of their probability of occurrence.

### Experimental setup

Experiments took place in a w×h × l = 3.0 × 3.0 × 3.6 m sound-attenuated and echo-free room and were performed in complete darkness (see Van Bentum et al., 2017; 2021, for more detailed descriptions of the setup). Sounds were presented from speakers (Minx Min 12, Cambridge Audio, United Kingdom) that were positioned all around the listener’s head and controlled via real-time processing units (RP2.1 Tucker-Davis Technologies, System 3 or TDT-3, for short). Head orientation was recorded with the magnetic search-coil technique (Robinson, 1963). To that end, listeners wore a lightweight plastic glasses frame (glasses removed) with a magnetic search coil attached to its center. A laser pointer was attached to this frame, which ensured the measurement of pure head-saccades, without the co-occurring saccadic eye-in-head movements of natural gaze shifts.

### Stimuli

Sounds were generated in MATLAB (version 2018b, The MathWorks, Inc., Natick, Massachusetts, United States) stored offline, and played back at a sampling rate of 48,828.125 Hz. Prior to the experiments, sound levels were calibrated for each stimulus type and speaker location to ensure equal-level presentation at the location of the listener’s head. Stimuli were broadband Gaussian white-noise bursts (bandpass filtered between 0.5 and 20 kHz), with durations of 200 ms and an onset/offset ramp of 5-ms sine/cosine-squared windows were used. All sounds were generated with renewed random seeds, effectively creating uncorrelated noise. Sounds were presented at equal levels, randomly chosen between 55-62 dBA.

### Sound locations

Sounds were presented in the horizontal plane between -80 and +80 deg azimuth. For each sound pair, relative eccentricity was characterized with respect to the sagittal midline, with one sound being the most eccentric (*T*_*E*_) and the other sound being the more central (*T*_*C*,_ Fig. 2A). Sounds could be presented within the same hemisphere, or in opposite hemispheres: *within-hemisphere* sounds were presented on the same side of the head (either to the left or to the right of the sagittal midline), *cross-hemisphere sounds* were presented on opposite sides of the head (one sound to the left, the other to the right of the sagittal midline, see Fig. 2A for schematic example). In total, 116 unique combinations of two targets were presented (Fig. 2B). The same speaker locations were used for both cross- and within hemisphere conditions (Fig. 2C), with the same within-hemisphere conditions being mirrored left and right of the sagittal midline, causing a surplus of this condition (Fig. 2C). Separation angles between speakers ranged from 10 to 60 deg in steps of 5 deg (Fig. 2D).

**Figure 2.**
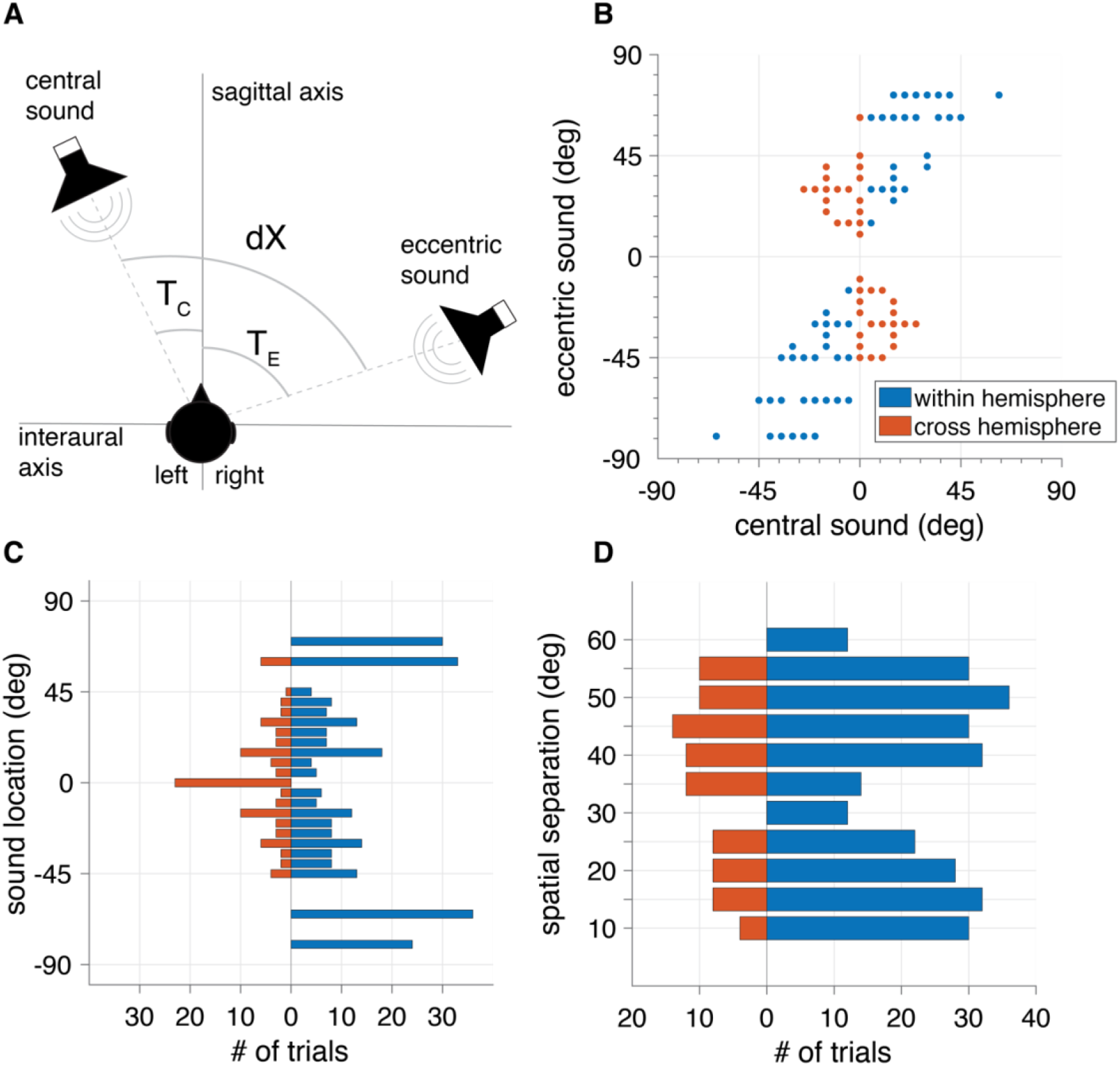
Experimental conditions. **(A)** Schematic top view of a listener in the experimental setup. Speaker pairs at central (T_c_, closest to mid-sagittal plane) and eccentric (T_e_, furthest from mid-sagittal plane). dX denotes spatial separation (|T_e_ – T_c_| in deg). **(B)** Locations of central-eccentric sound pairs. Each dot represents a unique combination of central and eccentric speaker locations in the experiment. Blue/red represent within/cross hemisphere condition, respectively. **(C)** Histogram of presented sound locations. Note that within hemisphere sound pairs are presented more often than cross-hemisphere pairs. **(D)** Histogram of spatial separations for the two hemisphere conditions. Due to technical limitations, the 30 and 60 deg separation could only be used for the within-hemisphere combinations.

### Control session

To acquire subject-specific localization parameters, all listeners completed a single-sound control session where they were asked to make only single responses to single targets (376 trials). White-noise sounds with the same duration and bandpass-filtered content as in the main two-sound experiment were presented randomly over all locations used. Sounds were presented at randomly roved levels of 58-65 dBA.

### Paradigm

Listeners were informed that there could be either one or two sound locations in each trial, without providing prior information on the number of sources and their probability of occurrence. In each trial, unbeknownst to listeners, two sounds were presented at each possible combination of location and spatial separation (as described above; Fig. 2B, D), all randomly interleaved across trials. Per session, each listener responded to 376 trials, which took about 20 minutes to complete.

During the experiment, listeners were asked to fix their gaze on a green LED at (0,0) degrees (center of vision), and to press a handheld button to initiate a trial. After the button press, there was a pause of 300-800 ms (drawn randomly from a uniform distribution), after which the LED was turned off, followed 200 ms later by the presentation of the sounds. This procedure was chosen to minimize the predictability of sound-onset timing and to exclude potential after-effects of head-fixation. Listeners were instructed to localize quickly and accurately with one or two head movements based on the number of sounds they could identify. When listeners perceived two sounds, they were instructed to briefly hold their gaze at the first response location (to allow for accurate localization endpoint detection), and from there make a second head movement to the other perceived sound location (see Fig. 3 for a visualization of this process). No explicit instructions were given to listeners as to which sound should be localized first (as sounds were always synchronous). After the final perceived sound was localized in the trial, listeners were instructed to hold their gaze at that location until the fixation light reappeared at (0,0) deg. For trials with one head movement, this meant maintaining gaze at the single perceived location, for trials where two sounds were perceived, this meant holding gaze at the second perceived location.

**Figure 3.**
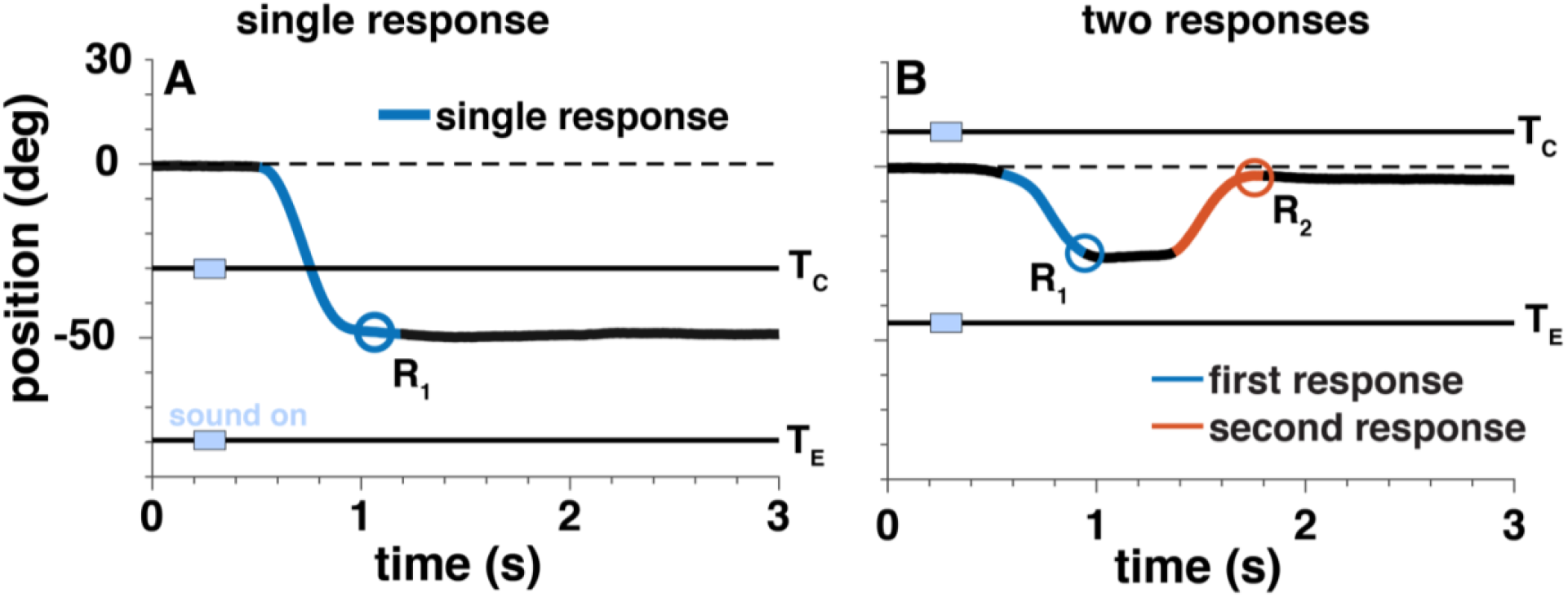
Example localization responses from listener L4. All responses started at the straight-ahead fixation point (0 deg; dashed line), were directed towards the perceived target(s), and maintained position until the trial ended (after 3 sec). Response endpoints R_1_ and R_2_ (as used in subsequent analyses) are indicated by the circles. Double-sound presentation is indicated by the light-blue bars. **(A)** Single response R_s_ = -50 deg directed in between T_C_ =-30 deg and T_E_ = -80 deg. Targets were presented within-hemisphere with a separation of 50 degrees. **(B)** Two responses were made to R_1_ =-25 deg and R_2_ =-2 deg for cross-hemisphere targets T_E_ =-45 deg and T_C_ = +10 deg with a separation of 55 deg.

### Data analysis

All data analysis was performed in MATLAB.

### Data selection

Using the procedure described above, coil signals were calibrated to azimuth and elevation angles for each measured sample. A custom-written MATLAB program was used to detect head saccades (e.g., Bremen et al, 2010). The threshold for automatic head-saccade onset- and offset detection was set at a head-velocity of 20 deg/s. Please note that this velocity was calculated from the two-dimensional movement; both azimuth and elevation components were considered. For example, responses that were made solely in elevation (even though all sounds were presented at 0 deg elevation) were included in the analysis and treated as a zero-azimuth response. Saccades were manually checked for irregularities (null-responses, anomalous profiles). Head-saccade endpoints were used for further analysis.

### Identification

Listeners were instructed to make one or two responses based on the number of targets they perceived. The ability to identify two stimuli at two different locations, regardless of localization accuracy, was estimated by taking the proportion of trials with two responses with respect to the total amount of trials for a given condition (pooled over listeners). To estimate the effect of spatial separation on sound identification, we fitted a generalized linear model with a logit link function on the binomially distributed data:

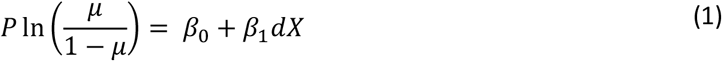

with *μ* ∈(0,1) the mean proportion of two responses, *β*_0_ and *β*_1_ predictor coefficients. Here, *β*_1_ is the predictor for spatial dependency and *β*_0_ a constant offset. In this model, each trial could have one or two responses based on a binomial distribution with parameter *μ* as the probability of observing two responses:

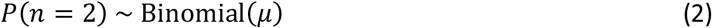

## Localization

### Single-target linear regression

To acquire subject-specific localization parameters, target-response regressions were performed on the single-sound control session, to quantify localization performance according to:

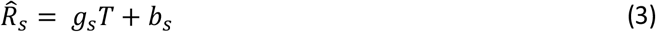

where 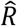 is the predicted response location (degrees azimuth), *T* is the target location (degrees azimuth), *g* is the regression slope (gain, dimensionless) and *b* is the offset (bias, in degrees), for listener s. The coefficient of determination (*r*^*2*^) between measured response and predicted response was also calculated. All listeners had gains close to 1.0 (between 0.96 and 1.2,), a negligible bias (between -0.2 and 8 deg) and r^2^ -values close to 1.0 (0.93-0.98), indicating highly accurate and precise localization performance. The linear gain and bias were used to correct for idiosyncratic listeners’ localization performance. To that end, the physical target locations (*T*) of the main two-sound experiment, were transformed to the response location predicted by the single-sound localization fit for each listener (Eqn. 3).

### Two-sound localization

In the two-sound experiment, listeners could make one (Fig. 3A) or two (Fig. 3B) responses in each trial. End point positions of the detected head saccades (i.e. at offset when head velocity fell below 20 deg/s) were taken as the measure of interest, both for single-response trials (Fig. 3A; *R*_*1*_) and two-response trials (Fig. 3B; *R*_*1*_, *R*_*2*_). In two-response trials, head movements were separated by a movement pause after the first response (Fig. 3B). Since there was no order in which participants were instructed to localize the sounds (these were presented synchronously and were perceptually identical), *R*_*1*_ and *R*_*2*_ could be directed to either target (central or eccentric). Note that for the execution of the head movement from *R*_*1*_ *to R*_*2*_ also required a coordination transformation to account for the new head-orientation at *R*_*1*_ after stimulus presentation.

Figure 3 shows a typical example of both a single and a two-response trial of one participant. In the single-response trial (Fig. 3A), the two sounds were presented in the listener’s left hemifield (within-hemisphere) at -30 and -80 deg (spatial separation is 50 deg). The participant made a single goal-directed head movement toward -50 deg with a reaction time of about 300 ms. In the two-response trial (Fig. 3B), targets were presented at T_E_ = -45 deg and T_C_ = +10 deg (thus, cross-hemisphere presentation with a separation of 55 deg). The first, leftward response ended at R_1_=-25 deg, and the second, rightward, response was initiated about 400 ms after the offset of R_1_, ending close to R_2_ = 0. Note that the end-points of the head movements always include some variability, meaning that from these single trials it is impossible to deduce whether the responses are goal-directed, directed to an average location, or to a location unrelated to the targets. We assessed this over the entire pool of trials, by determining the root-mean-square error between response and target locations. This was done for central, eccentric, and average (T_a_ = (T_e_ + T_c_)/2) target locations.

### Interaural correlation

To assess the influence of spatial separation and hemisphere presentation on sound identification and localization, we calculated the interaural correlation for two simulated broadband noise sounds. Sounds were presented at locations between ±80 deg azimuth and had a sound pressure level of 60 dB SPL. Simulated sound pressure *S*_*L*_*(t)* at the left ear and *S*_*R*_*(t)* at the right ear were calculated as follows:

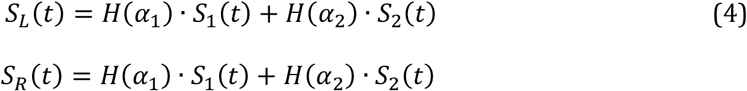

with the head-shadow attenuation function H(α_k_) = 9.7 · sin(0.02 · α_*k*_), with α_*k*_ the azimuth angle (deg) of sound k=1,2, and uncorrelated white-noise sound bursts, *S*_*1*_*(t)* and *S*_*2*_*(t)* (see also Van Wanrooij & Van Opstal, 2004). The interaural correlation was then calculated by taking the maximum of the cross-correlation between *S*_*L*_*(t)* and *S*_*R*_*(t)* over the full timeseries of the sounds (duration, *D* = 150 ms):

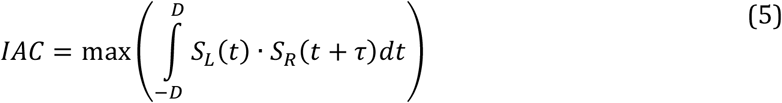

## Results

### Identification

To estimate the listener’s ability to identify two sources, we determined the relative proportion of trials with two responses. When two sounds were presented, identification performance of the listeners increased with source separation (Fig. 4). For small (<30 deg) separations, the proportion of two-response trials was below 0.5. Cross-hemisphere sounds evoked two responses far more often than within-hemisphere sounds (Fig. 4, red vs. blue) for all target separations. For the largest source separation with cross-hemisphere presentation (60 deg), participants perceived two sounds in more than 95% of all trials. In contrast, for the same separation, only 25% of trials with within-hemisphere presentation were perceived as two distinct locations. A GLM fit (see Methods; Eqn. 1) between spatial separation and proportion of two responses yielded offsets and slopes: *β*_0_ = −1.81 (95% CI: -2.4, -1.3), *β*_1_ = 0.06 (95% CI: 0.05, 0.08) for cross-hemifield presentation; and: *β*_0_ = −2.5 (95% CI: -2.9, -2.2), *β*_1_ = 0.03 (95% CI: 0.02, 0.04) for within-hemifield presentation. This entails that two responses occur in 50% of the trials at a spatial separation of ∼30 deg for cross-hemisphere presentation and of ∼80 deg for within-hemisphere presentation (i.e. outside of the range of spatial separations presented in this study).

**Figure 4:**
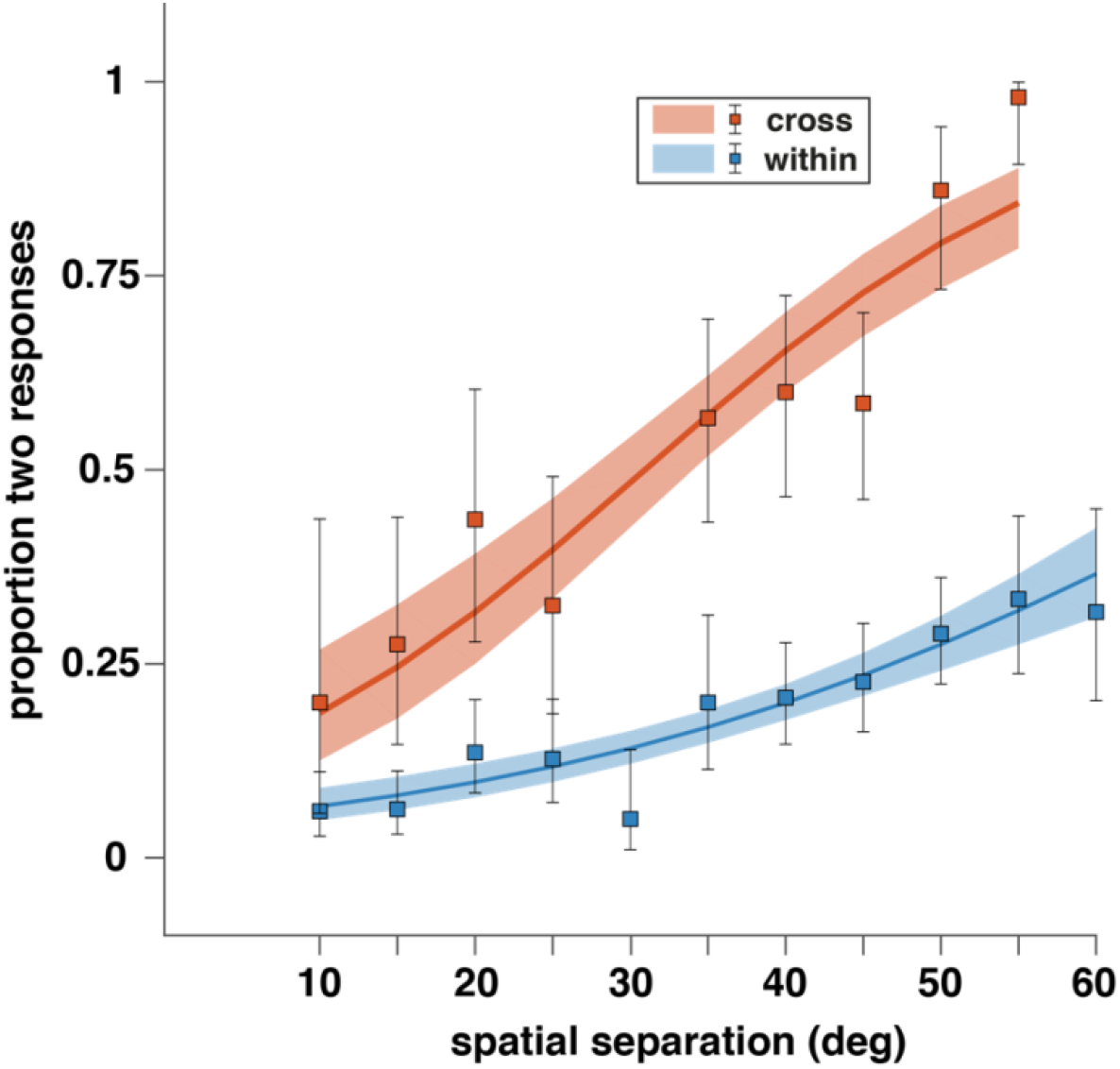
Source identification improves with source separation and cross-hemisphere presentation. Data points indicate proportion of trials with two responses, with 95% binomial confidence intervals as error bars. Solid lines indicate GLM model fit (Eqn. 1) over pooled data from all participants. Shaded areas indicate 95% confidence intervals for the fit. Red: cross-hemisphere sounds; blue: within-hemisphere sounds.

### Localization

We next analyzed the localization accuracy of the responses. According to instruction, listeners made one or two head movements in succession, briefly pausing at the first perceived location, and from there reoriented towards the second perceived location. Since the probability of making two responses increased with spatial separation (Fig. 4), especially when sounds were presented in different hemispheres, a different number of trials with one or two responses were obtained per unique (hemisphere, separation) condition.

#### Single and first responses are directed towards average sound location

To illustrate the analysis of the responses across trials for a single listener, Fig. 5 shows the response distributions of listener L1 as a function of three relevant stimulus variables: the perceived location (see Eqn. 3) of the central target, *T*_*C*_ (Fig. 5A,D), of the eccentric target, *T*_*E*_ (Fig. 5B,E), or of the average target, *T*_*avg*_ (Fig. 5C,F). The average-location target provided the best descriptor for both the single (top row) and the first (bottom row) head-movement responses. This can also be observed by the root-mean-square error between each target predictor and response (E), which was smallest for the average-location target, around 6-8 deg (Fig. 5C, F). In contrast, using central and eccentric sound locations as predictors gave much larger errors, between 18-23 degrees (Fig. 5A,B,D,E). The root-mean-square errors were near-identical for the within- and cross-hemisphere conditions (Fig. 5, blue and red, respectively), indicating that this listener’s sound localization was independent of hemisphere presentation (in stark contrast to sound identification for the entire group, cf. Fig. 4).

**Figure 5.**
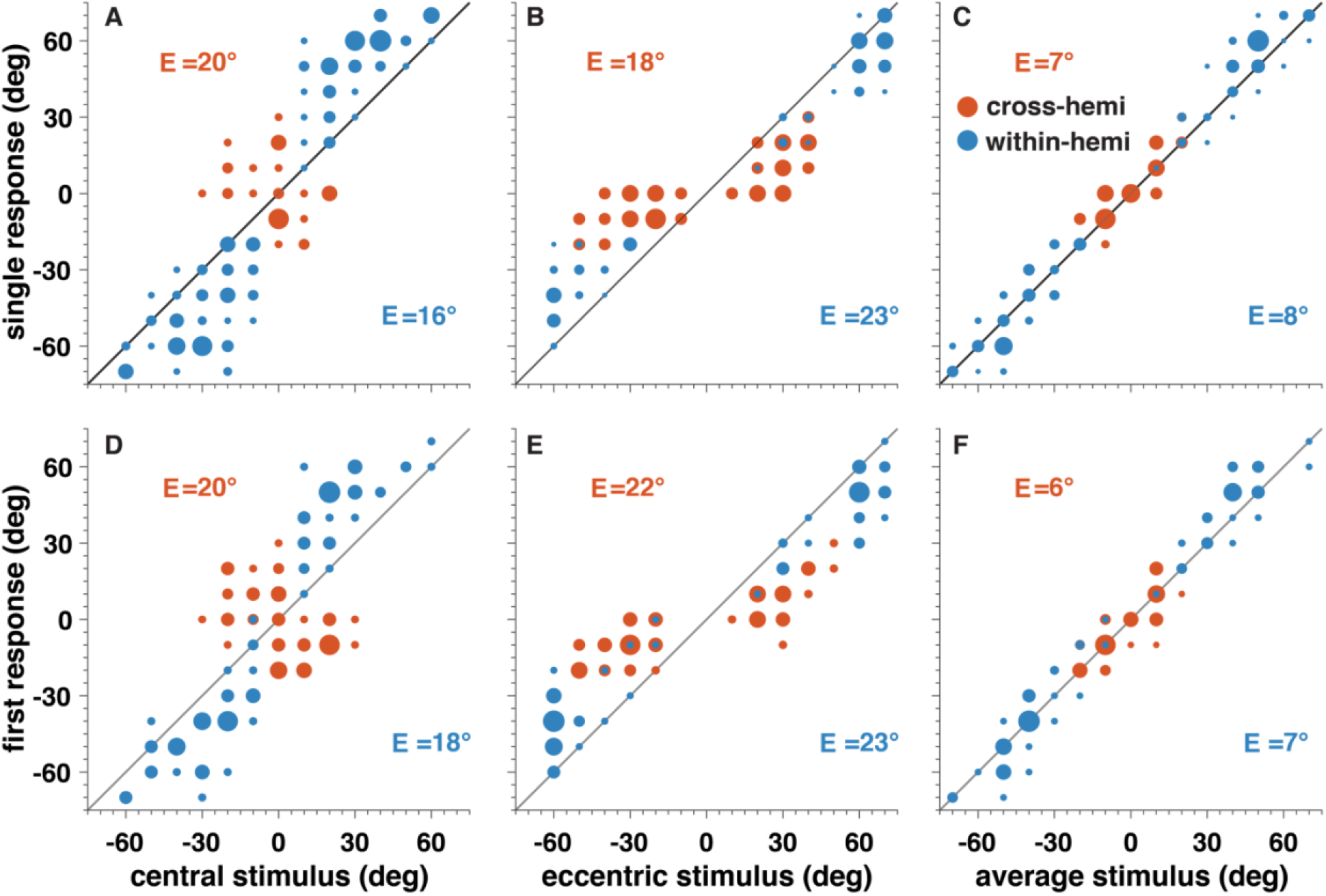
Single listener data shows averaging behavior in localization. Stimulus-response plots of listener L1 for single-response (top row) and the first of a double-response pair (bottom row) for three predictors. Errors E indicate root-mean-squared error of the regression, for within (blue) and cross (red) hemisphere conditions. **(A-C)** single response, **(D-F)** first of two responses as a function of the central **(A**,**D)**, eccentric **(B**,**E)** or average **(C**,**F)** sound location. Data are pooled across spatial separations, dX, but shown separately by different colors for the two hemisphere conditions. Size of the data points represents the relative number of responses. The highest correspondence to the response is obtained for the average-location stimulus for all situations **(C**,**F)**.

We quantified this for all listeners by calculating the root-mean-square error between response and perceived eccentric, central or average sound location (Fig. 6), for all spatial separations and both hemisphere presentations. For nearly all conditions, the average-location target was the best predictor, yielding the smallest error (Fig. 6, purple symbols vs orange and green). Errors increased with increasing spatial separation, in an almost linear fashion (Fig. 6, linear fit indicated by line and shaded areas). This increase was steepest for the eccentric- and central-target predictions (Fig. 6, orange and green).

**Figure 6.**
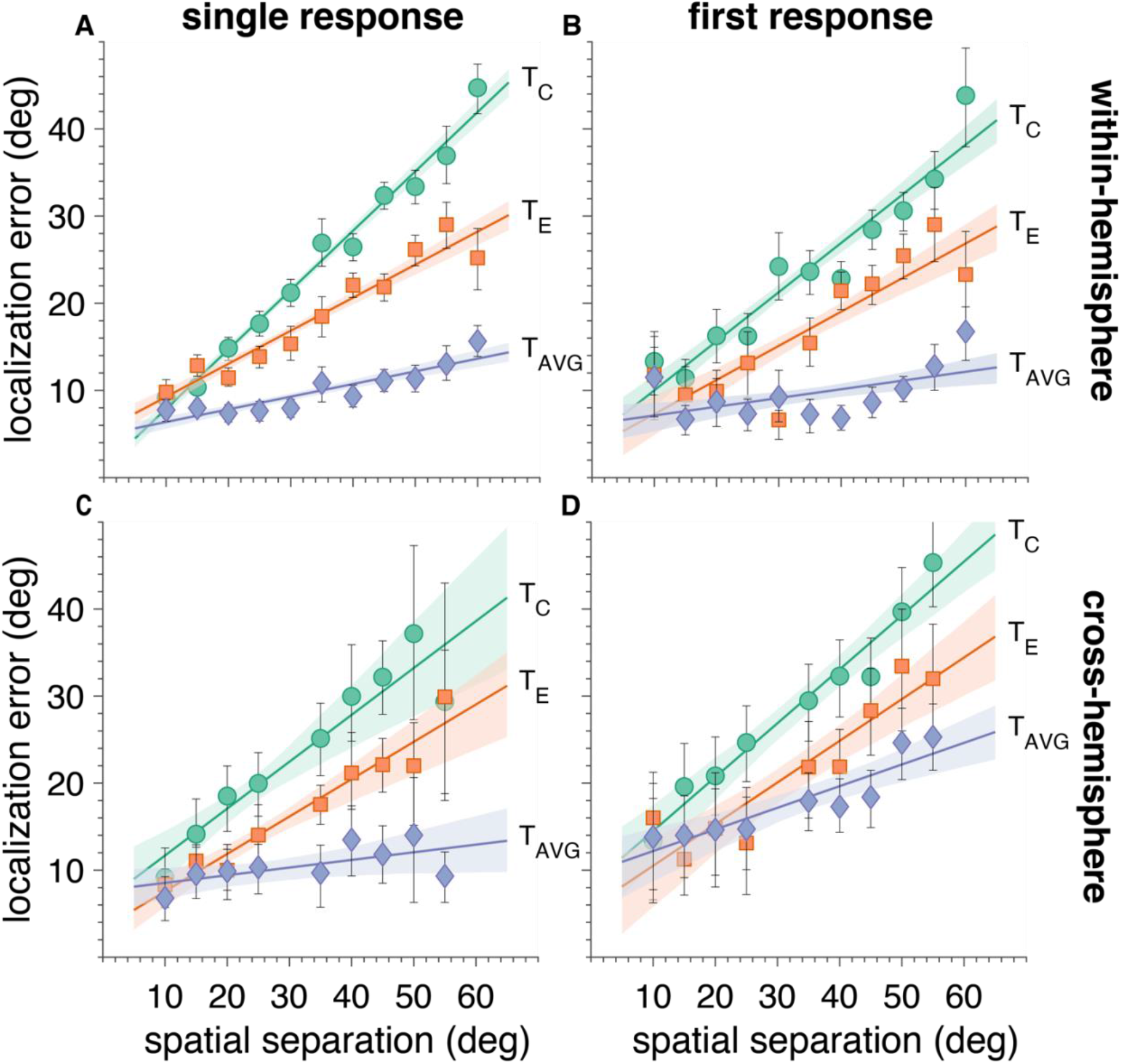
Averaging behavior in localization is consistent over stimulus conditions. Root mean-squared errors for the single **(A**,**B)** or first **(C**,**D)** response with respect to the central target (T_C_, green), the eccentric target (TE, red), and the stimulus average location (TAVG, blue), as a function of spatial separation (data pooled over listeners). Dots and error bars denote mean and standard deviation across listeners. **(A**,**C)** within-hemisphere sounds. **(B**,**D)** cross-hemisphere sounds. The smallest errors were obtained when the data were regressed against the average-location predictor for both the cross-hemisphere and within-hemisphere configurations.

### Second responses are not goal-directed

Second responses did not appear to be goal directed (Fig. 7). Instead, responses either returned towards the straight-ahead location estimate (‘central’ responses) or the head was displaced very little (‘null’ responses). To quantify this, we clustered the data in three groups using a k-means clustering analysis (Fig. 7A) according to how much the 2^nd^ response displaced the head away from the 1st-response endpoint (head shift *ΔH = R*_*2*_ *-R*_*1*_). We labelled the three response clusters, that were identified: the ‘null’ cluster, for which the head shift was negligible and independent of the first response (Fig. 7A, red); the ‘central’ cluster, for which the head shift seemed to be on average the same magnitude but opposite in sign of the first response, indicating that the second responses returned gaze to straight-ahead (Fig. 7A, blue); and a ‘ambiguous’ cluster, for which the data resembled either of the other two clusters, as both the first-response end points and the head shifts were around zero (Fig. 7A, green). In the target-response plots (Fig. 7B-G), the null-cluster were closer to the average-target location (Fig. 7D, red), than to the central (Fig. 7B) or eccentric (Fig. 7C) target location. Since the first responses were also close to the average-target location (Figs. 5, 6), this meant that the horizontal head shift of these second responses was small. The 2^nd^-response end-points of the central cluster, on the other hand, were not related to any of the predictors (Fig. 7E-G).

**Figure 7:**
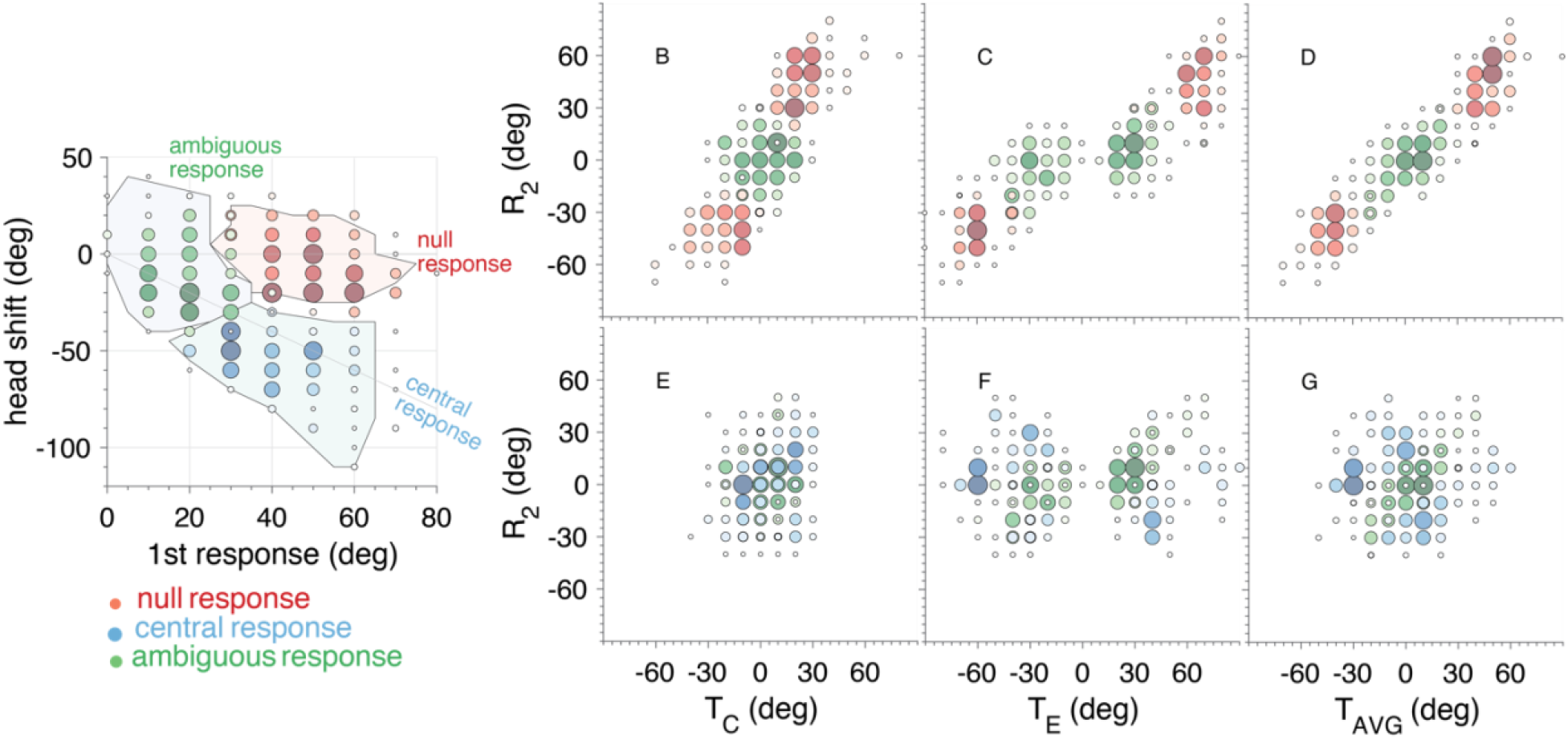
Second responses are not goal-directed. (**A)** Cluster analysis. Head shift ΔR (R_2_-R_1_) versus first response amplitude |R_1_|. Responses are pooled over listeners and stimulus conditions, and clustered into three groups: central (blue), null (red) and ambiguous (green). (**B-G**) Stimulus-response plots for second responses R_2_ after clustering. R_2_ versus central sound (**B, E**), eccentric sound (**C, F**) and average sound location (**D, G**). Null responses (B, C, D, red) display averaging behavior, where the second response stays close to the first (averaging) response location (see A). Central responses (E, F, G, blue) do not show a clear target-response relation; instead, they were oriented towards the estimated straight-ahead location (see A). Size of the data points represents relative number of responses.

## Discussion

### Summary

Listeners responded with goal-directed head movements to two uncorrelated synchronous broadband noise bursts, while horizontal spatial separation and hemispheric presentation was varied. They made one or two head movements based on the number of perceived sounds. Our results show that, while it was possible for participants to identify the presence of two sources, accurate localization of either source was impossible. Instead, listeners consistently responded to the average of the two target locations.

### Spatial segregation cues allow identification, but not localization

In the auditory system, temporal asynchrony is a strong cue for source segregation. Listeners can already identify and localize sounds in the horizontal plane with an onset delay of only a few milliseconds (Seeber and Hafter 2011; Brown et al., 2015; Van Bentum et al., 2021). Here, we focused on synchronous sounds, to eliminate temporal asynchrony as a possible segregation cue, leaving spatial separation as the only cue available (e.g., Best et al., 2004). Our results show that listeners were able to extract valid acoustic information arising from cross-hemispheric presentation and spatial separation to successfully identify the presence of two sounds (Fig. 4). In contrast, localization behavior consisted invariably of target averaging (Figs. 5-7), indicating that the segregation cues that helped to improve source-identification did not benefit sound-localization performance.

A potential cue that could allow listeners to identify the number of sounds, may be based on a broadband interaural-correlation mechanism (Faller and Merimaa, 2004; Dietz et al., 2011). Earlier studies have shown that a low interaural correlation can improve speech perception in noise and benefit spatial release from masking (May et al., 2013; Litovsky, 2012). To analyze whether our identification and/or localization data could be explained by such a mechanism (Fig. 8), we simulated the interaural correlation for any target-pair of synchronous, uncorrelated broadband noises in the frontal hemifield (Fig. 8A; Methods, Eqns. 4 and 5) and interpolated the identification probability (Fig. 8B; cf. Fig. 4) and localization variability (Fig. 8C; cf. Fig. 6) to these target pairs. A combination of large spatial source separation and cross-hemispheric presentation leads to low interaural correlation (data in the upper-left and lower-right quadrants), whereas close spatial proximity and within-hemispheric presentation produces a high interaural correlation for the (uncorrelated) noise bursts (the main diagonal). Identification accuracy (Fig. 8B; taken from Fig. 4) was highest for target combinations that yielded the lowest interaural correlation (cf. Fig. 8A). Overall, the identification pattern strongly resembled the interaural-correlation pattern. We propose that this strong negative correlation is not spurious, and that free-field source identification might indeed rely on the interaural correlation.

**Figure 8.**
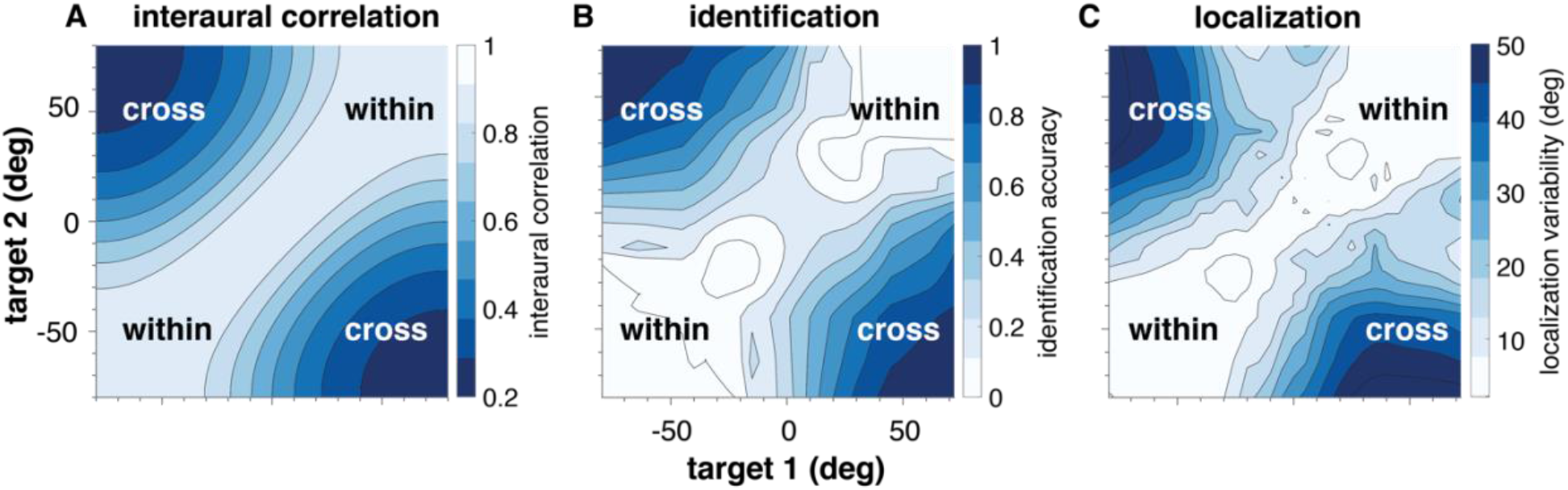
Target-pair locations giving rise to **(A)** interaural correlation, **(B)** identification accuracy, and **(C)** localization variability. (**A**) Interaural correlation between two uncorrelated broadband sounds presented at different locations in azimuth (Methods, Eqns. 4 and 5). Within- and cross-hemisphere sound pairs correspond to the different quadrants. (**B**) Identification probability and (**C**) root-mean-square error with respect to the average-target predictor (from the same data as shown in Fig. 4 and Fig. 6), both linearly interpolated for each target pair as in panel (A). Blue-white color scales indicate the magnitude of each measure.

Our (mean) localization data (Figs. 5, 7) does not immediately seem to depend on spatial separation or on hemisphere presentation. However, the root-mean-square error (Fig. 6) does seem to be larger for increasing spatial separation and for cross-hemisphere presentation. Taking this error (with respect to the average-location predictor) as a measure of response variability (Fig. 8C; taken from Fig. 7; note that the root-mean-square error equals the standard deviation of the residuals), a pattern emerges that corresponds nicely to the interaural-correlation map (cf. Fig. 8A), with low interaural correlation coinciding with large localization errors. We hypothesize that listeners could have perceived a broader spatial blur for the acoustic event when the interaural correlation was low (suggesting the presence of multiple sources at different locations) than when the interaural correlation was high (which would be more in line with a single point source). In principle, this might yield a potential cue to aid in localization of two synchronous sounds as the spatial blur extent may be informative on the target-pair locations (as discussed later in the Internal Priors section). Still, none of the listeners in the current experiment seemed to be able to use such a strategy to localize.

### Persistent spatial averaging

Our data demonstrate consistent spatial averaging, for single responses, and for first responses (Figs. 5-6). This finding corroborates and extends earlier studies on localization of horizontally and vertically separated synchronous sounds (Blauert, 1971; Perrot et al., 1987; Brown et al., 2015; Seeber and Hafter, 2011; Van Bentum et al., 2021; Bremen et al., 2010; Van Bentum et al., 2017; Ege et al., 2018). Secondary responses were never goal-directed; not to central, eccentric, or average target locations. Instead, these were either directed towards the estimated straight ahead or they did not move appreciably in the horizontal direction, thus staying near the perceived average azimuth (Fig. 7).

In the visuomotor system, saccadic eye-movement studies have shown that accurate localization and identification of synchronously presented visual targets is possible, given sufficient (>60 deg) retinal separation of the targets (Findlay, 1982; Ottes et al., 1984; Van Opstal and Van Gisbergen, 1990; Van der Stigchel and Nijboer, 2013). However, for relatively small target separations (up to about 45 deg) the visuomotor system invariably produces target-averaging responses (Findlay, 1982; Ottes et al. 1984; Van der Stigchel and Nijboer, 2013). This so-called ‘global effect’ (Findlay, 1982) has been explained by spatial interactions within the saccadic motor map of the midbrain Superior Colliculus (SC; Ottes et al., 1984; Van Opstal and Van Gisbergen, 1989). Evidence for averaging in the saccade-related responses of SC neurons has been provided by single-unit recordings in the monkey during a double-target response task (Van Opstal and Van Gisbergen, 1990). So far, similar explicit neural evidence for spatial averaging in the auditory system is lacking, although monkey SC neurons have been shown to also mediate eye-centered audio-motor responses, just like in the visuomotor system (Jay and Sparks, 1984). We recently postulated a conceptual model that could explain the occurrence of spatial averaging for synchronous and asynchronous sounds based on spatial-temporal interactions among neurons in the SC motor map (Van Bentum et al., 2021). The single-response and first-response data reported in the present experiments can be explained by such a mechanism as well.

### Internal priors

Due to the very low probability in natural environments that multiple sounds arising from independent objects produce exactly synchronous onsets, the auditory system will have developed a strong internal prior that synchronous cues, even when ambiguous, most likely originate from a single source and hence from a single location. Forcing the brain to accept that synchronous sounds were produced by more than one source may require strong additional prior information, such as instruction, added visual information, or allowing listeners to use dynamic cues induced by self-motion.

Yost and Brown (2013) demonstrated that with explicit instructions and repeated presentation listeners were able to correctly indicate the location of two uncorrelated, synchronous sounds. Listeners knew that there would always be two sounds, which imposed a cognitive prior on their auditory system, which could effectively bypass the identification stage (Fig. 1). Santala and Pulkki (2011) had also found that with repeated stimulus presentation listeners could identify up to three sound locations in the azimuth plane, possibly by making use of dynamic head-movement cues.

It is conceivable that because of the additional non-acoustic prior knowledge provided in these experiments, listeners may have followed a different response strategy than in our experiments, in which participants had to respond with fast and accurate head movements and lacked any prior information about the number of sounds. Moreover, sound presentation was too short to make use of potential additional dynamic head-movement cues. For example, as Fig. 8C indicates, the summed auditory event of a double-sound stimulus appears to induce a larger spatial extent than that produced by a single sound. Listeners could thus have used this information by responding to either side of the perceived auditory event in consecutive identical trials (or while probing with their head), thereby leading to reasonably accurate location estimates of the targets. To test for this interesting possibility, prior information should be systematically manipulated, e.g. by varying the (perceived) probability of the number of sound sources.

### Conclusion

Improved identification performance was unrelated to the performance in localization. This strongly suggests that the putative ‘what’ and ‘where’ pathways in the auditory system may be uncoupled (Fig. 1). We further conjecture that the ‘what’ pathway of the human auditory system can employ an interaural-correlation mechanism to identify the presence of multiple synchronous sources in the scene. This may be particularly useful in situations where a target signal needs to be segregated from ambient background noise. In contrast, the ‘where’ pathway appears to assume that synchronous sounds always originate from a single source location, which results in localization responses that are invariably directed towards the average target.

